# Integrated Multi-Omics Identifies Lineage-Dependent Myeloid Cells Recruitment and the APP-CD74 Axis as an Immunoregulatory Target in Pediatric High-Grade Glioma

**DOI:** 10.64898/2026.05.01.722277

**Authors:** Zhuang Wang, Ambuj Kumar, Banlanjo Umaru, Archana Mahadevan Iyer, Komal Khan, Hao-Han Pang, Maryam Fouladi, Rachid Drissi

**Author notes:** Correspondence: Rachid Drissi, Center for Childhood Cancer Research, Nationwide Children’s Hospital, 575 Children’s Crossroad, Columbus, OH.

## Abstract

Diffuse intrinsic pontine glioma (DIPG) is a devastating pediatric brain tumor with limited treatment options. Emerging evidence indicates infiltration of tumor-associated macrophages (TAMs) within tumor sites, accompanied by an immunosuppressive tumor microenvironment (TME). However, the mechanisms underlying macrophage recruitment and communication between TAMs and other cellular compartments within brain tumors remain poorly understood. Bulk RNA sequencing of 26 DIPG autopsy specimens with matched normal brain tissue, single-cell RNA sequencing data from eight DIPG patients integrated with public pediatric high-grade glioma (pHGG) datasets, and *in vitro* transwell and flow cytometry assays collectively indicated that DIPG tumors actively recruit monocytes through chemokine-mediated mechanisms. The chemokine expression of tumor cells is driven by a mesenchymal-like (MES-like) lineage state rather than histone mutations, as evidenced by significant correlation between MES-like lineage scores and chemokine expression scores across 46 pHGG cell lines. CellChat analysis identified APP–CD74 signaling as a prominent tumor cell-TAM interaction pathway, supported by immunofluorescence validation. Notably, *APP* expression was significantly reduced in DIPG tumor tissues compared with normal brain tissue at both the RNA and protein levels. Recombinant APP stimulation of THP-1–derived macrophages induced a robust proinflammatory response, including upregulation of M1-like markers, enrichment of interferon-related pathways, and elevated secretion of inflammatory cytokines. Collectively, these findings indicate that *APP* suppression in tumors attenuates the antitumor activity of TAMs and promotes an immunosuppressive microenvironment. Furthermore, protein modeling and docking analyses identified the APP-CD74 binding interface, providing a structural basis for therapeutic targeting.

## INTRODUCTION

Pediatric high-grade gliomas (pHGG) are rare but uniformly aggressive and fatal central nervous system (CNS) tumors with multiple subtypes [50]. Diffuse intrinsic pontine glioma (DIPG) is the most aggressive subtype of pHGG, arising in the pons and characterized by diffuse infiltrative growth, rapid progression, and a dismal prognosis with a median overall survival of less than 12 months [9]. A defining hallmark of DIPG is the lysine-to-methionine substitution at histone H3 lysine 27 (H3K27M), which disrupts epigenetic regulation and drives oncogenic transcriptional programs [36, 78]. Due to the anatomical location, infiltrative nature, and protective effect of the blood-brain-barrier (BBB), the current standard of care remains limited to palliative radiotherapy [17]. Recently, Modeyso (ONC201) was approved by the Food and Drug Administration (FDA) for the treatment of diffuse midline glioma (DMG) with H3K27M mutations, expanding therapeutic options for this devastating disease [58]. Additionally, multiple clinical trials are evaluating chimeric antigen receptor T (CAR-T) cell therapies targeting various antigens, such as disialoganglioside GD2, B7 homolog 3 (B7H3), and Interleukin-13 receptor subunit α2 (IL13-Rα2), for pediatric high-grade glioma (pHGG), including DIPG and DMG [3, 4, 26, 57, 72]. These studies underline the promising potential of CAR-T therapies for pediatric brain tumors while highlighting critical challenges, including efficient delivery and the toxicity associated with cytokine release syndrome (CRS). Beyond tumor cell–intrinsic factors, the tumor microenvironment (TME) has a crucial role in disease progression and therapeutic resistance in pediatric brain tumors, including DIPG [60]. Emerging high-throughput technologies, such as single-cell RNA sequencing and spatial transcriptomics, have enabled high-resolution characterization of the TME, providing insight into the molecular programs of both tumor cells and infiltrating immune populations. Brain tumor cells, including those in DIPG, have been shown to display lineage-specific states, including neural progenitor-like (NPC-like), mesenchymal-like (MES-like), oligodendrocyte precursor-like (OPC-like), and astrocyte-like (AC-like) states [25, 34, 48]. These states are, in part, governed by the H3K27M-driven epigenetic reprogramming, and tumor cells retain the plasticity to transition between states, which promotes intratumoral heterogeneity and therapeutic resistance. In contrast to adult glioblastoma multiforme (GBM), DIPG is characterized by sparse infiltration of lymphocytes (T and B cells) and a predominance of myeloid populations, including microglia and monocyte-derived macrophages [46, 56]. This immunosuppressive TME of DIPG poses major challenges for the development and efficacy of current immunotherapies. Although recent studies have begun to delineate the immune landscape of DIPG, the contributions of specific immune cell subsets to tumor pathogenesis remain poorly understood. In particular, the interactions between tumor cells and tumor-associated macrophages/microglia (TAMs) are not well defined [35, 71]. Addressing these knowledge gaps is critical for reprogramming the TME toward a pro-inflammatory state and for identifying novel therapeutic targets.

## MATERIALS AND METHODS

### Clinical cohort

This study was approved by the Institutional Review Board (IRB) at Nationwide Children’s Hospital (NCH; IRB: 2016-3357) and Cincinnati Children’s Hospital (CCHMC, IRB:2019-1220). It included patients enrolled in the Pediatric Brain Tumor Repository (PBTR), a multidisciplinary approach of autopsy donation from pediatric brain tumor patients [28, 85]. Tumor specimens and matched normal frontal lobe tissue were obtained at autopsy under an IRB-approved protocol after written informed consent was obtained from patients and their families. All clinical data were de-identified prior to analysis to ensure patient confidentiality and adherence to ethical standards.

### Cell lines and cell culture

CCHMC-DIPG-1 (H3WT) and CCHMC-DIPG-2 (H3.3K27M) are primary patient-derived DIPG cell lines established in our laboratory and authenticated by whole-genome sequencing and RNA sequencing, as previously described [65]. SU-DIPG-IV (H3.1K27M) and SU-DIPG-XXXVI (H3.1K27M) were kindly provided and authenticated by Dr. Michelle Monje (Stanford University). All DIPG cell lines were maintained according to established laboratory protocols [42]. THP-1 cells were purchased from American Type Culture Collection (TIB-202, ATCC, Manassas, Virginia, USA) and validated by the supplier using short tandem repeat (STR) profiling. THP-1 cells were cultured in RPMI-1640 medium (30-2001, ATCC) supplemented with 10% fetal bovine serum and 1% penicillin-streptomycin at 37°C, 5% CO_2_. All cell lines were confirmed to be free of mycoplasma contamination prior to experimentation.

### Transwell migration assay

The transwell migration assays were performed using 24-well inserts with 3-μm pore-size PET translucent membranes (9023002, cellQART, Grand Rapids, MI, USA). THP-1 cells were stained with 10 μM CellTracker™ Orange CMRA (C34551, Thermo Fisher) for 30 minutes at 37°C according to the manufacturer’s procedure. A total of 2.5×10^4^ THP-1 cells were seeded into the upper chamber of each insert, while the lower chamber was filled with 5×10^4^ DIPG cells or conditioned media. Cell migration was monitored using the Incucyte SX5 Live-cell Imaging system (Sartorius). Fluorescent-labeled THP-1 cells that migrated to the lower chamber were detected and quantified every 2 hours. For each well, four fields of view were captured and the average number of fluorescent-labeled cells per field was calculated to represent the final number of cells migrated to the lower chamber.

### Flow cytometry

Flow cytometry was performed to assess the phenotype of THP-1 cells after coculture with DIPG cells. Briefly, single cells were washed twice with phosphate-buffered saline (PBS) and resuspended in FACS buffer (PBS with 2% fetal bovine serum and 2mM EDTA). Fc receptors were blocked by human FcR blocking antibody (14-9161-73, Invitrogen) for 15 minutes at 4°C. Then, cells were stained with an antibody cocktail consisting of CD45-PE (clone 2D1, Invitrogen), CD11b-BV421 (clone M1/70, BioLegend), CD163-Red718 (clone MAC2-158, BD Biosciences), and HLA-DR/DP/DQ-FITC (clone YE2/36-HLK, Invitrogen) for 45 minutes at 4°C in dark. After staining, cells were washed twice and resuspended in FACS buffer for acquisition. Data were collected on Cytek Aurora Spectro flow cytometer and analyzed using FlowJo v10 (Tree Star, Ashland, OR, USA). Gating strategies were applied to identify CD45^+^ THP-1 cells and further quantification and analysis.

### Amyloid Precursor Protein (APP) treatment of THP-1-derived macrophages

THP-1 monocytes were differentiated into macrophages by plating 5×10^5^ THP-1 cells in 6-well plates and stimulating with 100ng/mL phorbol 12-myristate 13-acetate (PMA) for 48 hours. Then, cells were washed and rested in PMA-free culture medium for 24 hours. Subsequently, cells were stimulated with recombinant human APP751-his (HY-P704352, MedChemExpress) at a final concentration of 250ng/mL in complete RPMI medium for 8, 24, and 72 hours. Protein storage buffer of recombinant human APP was used as a vehicle control.

### Immunofluorescence

Immunofluorescence staining was performed as reported earlier [64]. Frozen tissue slides were incubated with primary antibodies against APP (E4H1U rabbit mAb, Cell Signaling Technologies, Danvers, MA, USA) and CD74 (VIC-Y1 mouse mAb, Invitrogen, Waltham, MA, USA) followed by staining of secondary antibodies staining with donkey anti-rabbit IgG-Alexa Flour568 and donkey anti-mouse IgG-Alexa Flour647 antibodies (Jackson ImmunoResearch Laboratories Inc., USA). Samples were mounted and stained with antifade mounting medium containing DAPI (H-1200-10, Vector Laboratories, Plain City, OH, USA) then scanned using Nikon Ti2 Confocal Microscope. Image processing was done with NIS-Elements AR (v5.42.03).

### Western blot

Western blot assays were performed following the established protocol described previously [69]. Briefly, the membranes were blocked by 3% non-fat milk and overnight incubation with the following primary antibodies: anti-APP (E4H1U, Cell Signaling Technologies, 1:1000) and anti-β-Actin (8H10D10, Cell Signaling Technologies, 1:1000). Then, the membranes were stained with corresponding HRP-conjugated secondary antibodies. The images were captured with ChemiDoc MP Imaging system and quantified band intensity with Image Lab version 6.1 (Bio-Rad Laboratories).

### Luminex-based multiplex cytokine assay

For the multiplex cytokine assay, 2.5×10^5^ DIPG cells were seeded per well and cultured for 24 hours under previously described culture condition. Cell culture supernatants were collected, centrifuged at 500×g for 5 minutes to remove debris, and submitted to Eve Technologies Corporation (Calgary, Canada) for analysis using the human Cytokine/Chemokine 71-plex Discovery assay^®^ Array (HD71) and TGFB 3-plex Discovery assay^®^ multi-species array. Cytokine/chemokine concentrations were quantified by comparison with standard curves generated for each analyte; values higher or lower than designated concentration values for specific analytes are defined as out of range (OOR).

### RNA sequencing

RNA extraction is performed with RNeasy Plus Mini Kit (Qiagen, Hilden, Germany) in accordance with the manufacturer’s protocol as described in previous work [85]. RNA concentration is quantified using Qubit RNA BR assay kit (Qiagen). Transcript abundances were quantified using Salmon (v1.11.0) with the GRCh38 reference transcriptome (GENCODE release 21). Quantification files were imported into R (v4.4.1) for downstream analysis. Briefly, the differential expression analysis was performed with the limma-voom pipeline (v3.54). For visualization, gene expression was summarized with TPM normalization, followed by a log2(x+1) transformation. Pathway enrichment analysis was conducted using clusterProfiler (v4.2.2) and GSEA with gene sets from the Molecular Signature Database (MSigDB, v2024.1.Hs). The p-values were adjusted for multiple testing using the Benjamini-Hochberg method.

### Transcriptional chemokine and lineage scores with ssGSEA

To evaluate the relation between chemokine expression and tumor cell lineage, single-sample gene set enrichment analysis (ssGSEA) was applied to the RNA-seq data of 46 pHGG cell lines obtained from Childhood Cancer Model Atlas (CCMA) [67]. ssGSEA was implemented using the GSVA package (v1.52.3) to calculate the enrichment score per cell line [23]. The chemokine gene set contains 9 chemokines with established roles in myeloid recruitment as follows: *CCL2, CCL3, CCL4, CCL5, CCL7, CXCL8, CXCL10, CXCL12*, and *CX3CL1*. The chemokine score computed based on this gene set reflects the transcriptional program of chemokine expression of each cell line. The lineage scores of 4 cell states, including OPC-like gene set (*PDGFRA, OLIG1, OLIG2, SOX10, GPR17, PLP1, VCAN*, and *OMG*); the NPC-like gene set (*SOX11, SOX4, DLL1, DLL3, HES6, ASCL1, TUBB3, STMN2, DCX*, and *CD24*); the MES-like gene set (*CHI3L1, CD44, SPP1, VIM, FN1, ANXA2, LGALS3*, and *TIMP1)*; and the AC-like gene set (*GFAP, S100B, SLC1A3, AQP4, SPARCL1, FABP7, METTL7B*, and *EDNRB*) were computed for each cell line based on marker genes adopted from Neftel et al. [59]. The dominant lineage identity was assigned to each cell line based on its highest lineage ssGSEA score. Pearson correlation coefficients were calculated between chemokine score and each lineage scores across all 46 pHGG cell lines with linear regression fits. Statistical significance was assessed by two-tailed t-distribution and statistical significance with p < 0.05.

### Single-cell RNA sequencing library preparation

Tumor specimens were processed and dissociated into single cells. The sequencing library preparation and sequencing were done at Cincinnati Children’s Hospital Medical Center Gene Expression Core using 10x Genomic Single Cell 3’ Gene Expression v3 chemistry (10X Genomics, Pleasanton, CA, USA), and sequencing was performed on the Illumina NovaSeq 6000 platform.

### Single-cell RNA sequencing analysis

Raw sequencing reads were demultiplexed and converted to FASTQ files, followed by alignment to the human reference genome (build GRCh38), barcode processing, UMI counting, and gene expression matrices generation using CellRanger (v4.0.0) on 10x Genomics Cloud Analysis. Count matrices were imported into R and filtered, normalized, and processed in Seurat (v4.0.0) [24]. Low-quality cells were excluded from analysis, and doublets were further identified and removed using DoubletFinder [53]. Subsequently, gene expression counts were normalized, scaled, and the top 3,000 highly variable features were identified with FindVariableFeatures [24]. To correct the batch effect of patient samples, we conducted data integration with the Harmony package and principal component analysis for downstream analysis [38]. Uniform Manifold Approximation and Projection (UMAP) was used for visualization [54]. Cell type annotation was performed by examining the expression of canonical markers of tumor cells, immune cells, and stromal cells. Marker genes and cell populations were validated with published analyses of GBMs and pHGG datasets [1, 48, 55]. The cell communication analysis was performed using CellChat with human CellChatDB ligand-receptor interaction database [32]. Visualizations of results were generated using built-in functions of CellChat, Seurat, and ggplot2 packages [76]. The analysis workflow is available at: https://github.com/Ambuj/UF/DrissiLab_Glioma_Single_Cell_Analysis.

### Protein modeling

Amino acid sequences for human amyloid precursor protein (APP; UniProt accession P05067) and CD74 (UniProt accession P04233) were obtained from UniProt [70]. Initial three-dimensional models of APP and CD74 were generated using I-TASSER-MTD, which combines threading-based template identification with iterative full-length structural assembly and refinement for multi-domain proteins [84]. The predicted monomeric models were energy-minimized in GROMACS to relieve steric clashes and optimize local geometry [2, 6]. Minimization was performed using the steepest descent algorithm with a maximum force convergence criterion of 1000 kJ mol^−1^ nm^−1^, a step size of 0.01 nm, and a maximum of 100,000 steps. Electrostatic interactions were treated using the particle mesh Ewald (PME) method, with 1.2 nm cutoffs for both Coulombic and van der Waals interactions [11]. Hydrogen bonds were constrained using the LINCS algorithm [2]. To construct the oligomeric assembly of CD74, the minimized monomeric CD74 model was structurally superposed onto the CD74-containing oligomeric arrangement observed in the cryo-EM structure of the human invariant chain complex with HLA-DR15 (PDB: 8VRW) using PyMOL [13]. The resulting oligomeric CD74 model was subsequently subjected to a second round of energy minimization in GROMACS before downstream docking analyses. For membrane-associated structural modeling, transmembrane segments in APP and CD74 were predicted using the membrane protein topology prediction method TMHMM-2.0 [39]. The resulting protein models were embedded into a lipid bilayer using CHARMM-GUI Membrane Builder, which was used to orient the transmembrane regions relative to the membrane normal and generate the protein-membrane system for subsequent structural refinement and analysis [33, 77]. Surface hydrophobic patches were identified using the Protein-Sol Patches server to map hydrophobic regions that could contribute to membrane interactions or intermolecular associations. Protein-protein docking between the CD74 oligomer and APP was then performed using HADDOCK [27, 29].

### Membrane Restrained Gaussian Network Model

Elastic network models (ENMs) provide a computationally optimized model for characterizing the intrinsic dynamics of protein structures. Although these models do not capture the full atomistic detail accessible through molecular dynamics simulations, they have been shown to reliably reproduce the dominant collective motions of proteins, particularly the low-frequency modes that are most strongly associated with biological function. These slow, large-scale conformational fluctuations are often difficult to sample adequately using conventional molecular dynamics due to timescale limitations. Among ENM approaches, the Gaussian network model (GNM) remains one of the earliest and most widely used methods that has been shown to correlate well with experimentally measured crystallographic B-factors [5, 40, 41]. Here, GNM was applied to characterize residue-level dynamics and interfacial mechanical coupling within the docked APP–CD74 complex. The CD74 oligomer and APP protein residues were represented as a coarse-grained elastic network with each residue modeled as a single node at the C*α* position. Pairs of residues *i* and *j* were connected by identical harmonic springs with force constant *γ* when the corresponding C*α*–C*α* distance satisfied *r*_*ij*_ ≤ *r*_*c*_, where *r*_*c*_ = 7.5Å. The resulting network topology is encoded in the Ν × Ν Kirchhoff matrix Γ, where Ν is the total number of amino acids in the APP–CD74 complex. The energy is calculated as,

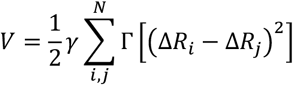

where Δ*R*_*i*_ and Δ*R*_*j*_ are the fluctuation vector of amino acids *i* and *j, γ* is the spring constant, and Γ is defined as

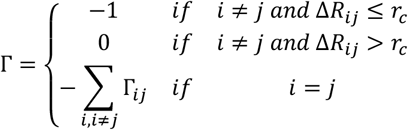

To incorporate the constraining effect of the membrane, amino acids in contact with lipid molecules in the docked protein–membrane assembly were identified using a 4.5 Å atom–atom distance cutoff and subjected to an additional harmonic restraint. Accordingly, the membrane-restrained Kirchhoff matrix was defined as

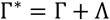

where,

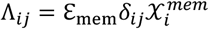

with *δ*_*ij*_ representing the Kronecker delta with 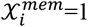 for membrane-contacting residues and 0 otherwise. The membrane restraint strength was set to ℇ_mem_ = 50. Intermolecular interface residues between CD74 and APP were identified independently from the docked complex using a 4.5 Å heavy atom contact distance criterion. Singular value decomposition of Γ^*^ was applied to obtain the lowest-frequency eigenvector (*ν*) and eigenvalues (*λ*). Top 10 non-rigid body eigenvectors and eigenvalues were used to calculate the pseudoinverse of Γ^*^ as,

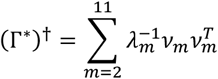

Residue fluctuations were calculated as,

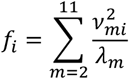

The covariance matrix was approximated from the pseudoinverse as shown below,

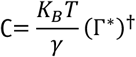

Interfacial mechanical coupling between contacting CD74–APP residue pairs was quantified using an effective stiffness metric,

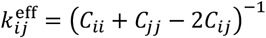

such that larger values correspond to smaller relative fluctuations and dynamic coupling. Because CD74 was modeled as a trimer, symmetry-equivalent residue positions from all three chains were consolidated by residue number and summarized against APP residues using the median 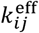.

### Statistical analysis

All statistical analysis of sequencing data was performed using the intrinsic methods implemented within the respective R packages. Downstream analysis, including differential gene expression and pathway enrichment, was carried out using default statistical frameworks embedded in each package. For experimental assays such as flow cytometry and cell migration, statistical analysis was conducted using GraphPad Prism (v10.6.0). Comparisons between multiple groups were analyzed by one-way or two-way analysis of variance (ANOVA) followed by appropriate post hoc tests. For comparison between two groups, we used unpaired two-tailed Student *t*-tests. Data presented as mean ± standard error of the mean (SEM), and p < 0.05 was considered statistically significant.

## RESULTS

### RNA sequencing reveals antigen presentation programs and myeloid activation in DIPG

To characterize the transcriptomic landscape of DIPG tumors, we performed RNA sequencing on 26 DIPG patient autopsy specimens along with paired normal brain tissues (left or right frontal lobes) **(Fig. 1a)**. As illustrated by the volcano plot, differential expression analysis revealed widespread transcriptomic alterations in the tumor tissues **(Fig. 1b)**. In total, 2352 genes were significantly upregulated, and 2458 genes were downregulated in the tumor cohort (|log2FC|>1, FDR<0.05). Notably, among the upregulated genes, there was a signature of immune modulatory factors, including multiple immune checkpoint genes (e.g., *PDCD1, LAG3*) and several key genes related to myeloid cells (e.g., *CSF1, CD163, CD86, CCL2*). Collectively, this transcriptomic profile suggests that DIPG tumors are characterized by increased macrophage infiltration and the establishment of a non-inflammatory immune microenvironment. To elucidate the biological functions of the differentially expressed genes, we performed Gene Ontology (GO) enrichment analysis on significantly upregulated genes in tumor samples using the clusterProfiler package, and the top enriched GO pathways were plotted **(Supp Fig. 1b-d)**. Among the top enriched pathways, five were related to MHC-mediated antigen presentation, underscoring the involvement of the immune system in TME **(Fig. 1c)**. Additionally, enrichment of pathways related to extracellular matrix organization, collagen binding, and integrin binding suggests activation of stromal remodeling processes that may facilitate tumor invasion. To better visualize the expression of genes contributing to these MHC-related pathways, we mapped individual genes into the gene-pathway network of the enriched MHC-associated GO terms. A broad induction of MHC class II protein complex genes (e.g., *HLA-DRA, HLA-DPA, HLA-DQA*), along with genes encoding associated chaperons, such as *CD74* and *HSP90AA1*, was observed **(Fig. 1d)**. Extending this analysis, we identified 15 significantly enriched MHC-related pathways in DIPG tumors, encompassing multiple biological processes associated with the MHC complex **(Supp Fig. 1a)**. This broad enrichment of antigen presentation pathways together with the upregulation of related genes, suggests that DIPG tumors retain immunological interfaces and active antigen presentation program, despite their well-recognized immunologically “cold” phenotype. Given the upregulation of myeloid genes in DIPG tumors, we next sought to investigate whether the expression of marker genes bears prognostic significance. The correlation analysis between overall survival and expression of marker genes for macrophages, including *ITGAM* (CD11b), *CD14*, and *STAT6* in tumor tissue, was conducted [10, 12, 81]. The higher expression of each of these genes was significantly correlated with shortened survival duration **(Fig. 1e)**. The data further support the role of tumor-associated myeloid cells in promoting disease progression and contributing to poor prognosis in DIPG.

**Figure 1.**
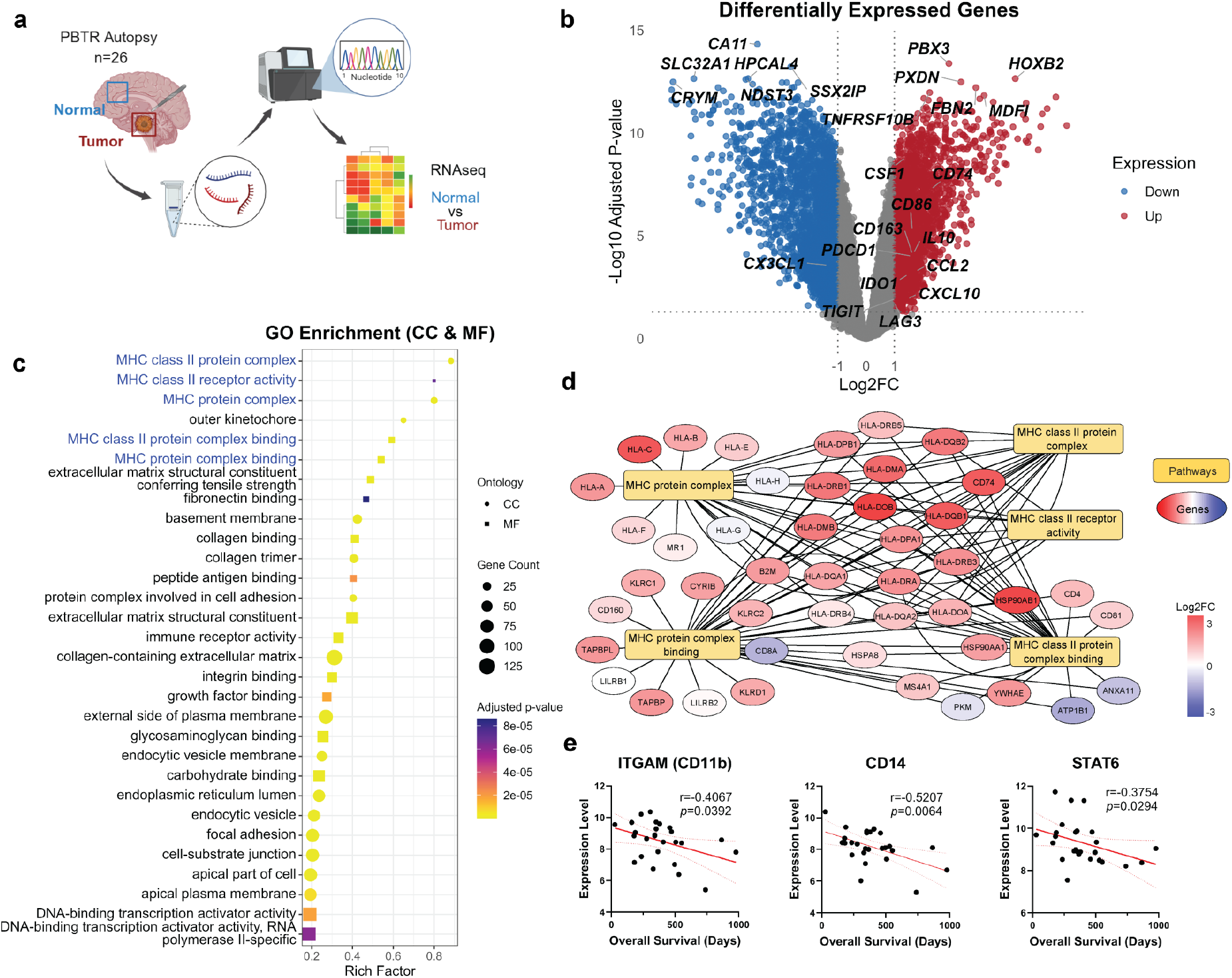
Bulk RNA-seq reveals immune and myeloid transcriptional signatures in DIPG. **(a)** Schematic overview of the RNA-sequencing workflow using post-mortem DIPG tumor and matched normal brain tissue (created with BioRender). **(b)** Volcano plot showing differentially expressed genes in DIPG tumor tissue compared to matched normal brain. Significantly upregulated genes are highlighted in red; downregulated genes are shown in blue. **(c)** Gene Ontology (GO) pathway enrichment analysis of significantly upregulated genes reveals enrichment in Major Histocompatibility Complex (MHC)-related pathways. **(d)** Network graph of individual genes contributing to the top MHC-related GO terms. The graph was generated by Cytoscape with pathways in yellow, upregulated genes in red, and downregulated genes in blue. **(e)** Correlation of myeloid-related gene expression (*CD11b, CD14, STAT6*) in tumor tissue with overall survival across DIPG patients.

### DIPG cells exhibit differential capacities to recruit and polarize monocytes

Having established the transcriptional upregulation of myeloid-related genes in DIPG tumors, we next examined monocyte and macrophage migration to elucidate the mechanisms by which tumor cells recruit immune cells into the TME. For this purpose, gene set enrichment analysis (GSEA) was performed to assess enrichment of Hallmark pathways in DIPG tumor samples [66]. We observed strong enrichment of positive regulation of *MCP-1* (*CCL2*) production, leukocyte migration, and macrophage activation, suggesting chemokine-driven monocyte recruitment and macrophage activation in DIPG tumors **(Fig. 2a)**. To gain a more detailed understanding of the chemokine profile driving myeloid cell recruitment in DIPG, we next examined the expression of individual chemokines and their corresponding receptors. A marked upregulation of *CCL2* was observed in tumor tissue, consistent with the GSEA results showing enrichment of positive regulation of *MCP-1* (*CCL2*) production. Additional chemokines, including *CCL3, CCL4, CXCL10*, and *CXCL12*, as well as chemokine receptors *CCR5* and *CXCR4*, were significantly upregulated in tumor samples **(Fig. 2b)**. Collectively, the enrichment of migration-related pathways and the increased expression of key chemokines and receptors underscore enhanced recruitment and retention of monocytes/macrophages within the DIPG TME. To validate findings from bulk RNA-sequencing data, we established an *in vitro* Transwell co-culture assay using human DIPG cells (CCHMC-DIPG-1, CCHMC-DIPG-2, SU-DIPG-IV, SU-DIPG-XXXVI) and human monocytic cell line (THP-1). DIPG cells or their conditioned media were seeded into the lower chamber of transwell migration system while THP-1 cells stained with CellTracker™ dye were seeded in the cell culture insert with 3µm pore membrane **(Fig. 2c)**. Surprisingly, THP-1 cells migration was markedly enhanced when co-cultured with SU-DIPG-IV (H3.1K27M) and SU-DIPG-XXXVI (H3.1K27M) cells or exposed to their conditioned media, whereas CCHMC-DIPG-1 (H3WT) and CCHMC-DIPG-2 (H3.3K27M) did not promote monocyte migration **(Fig. 2d, Supp. Fig. 2a)**. This differential capacity among DIPG cell lines was also observed in their ability to promote monocytes activation and polarization, as indicated by flow cytometry **(Fig. 2e and f)**. THP-1 cells cocultured with SU-DIPG-IV and SU-DIPG-XXXVI cells or treated with their conditioned media showed significantly increased proportions of immunosuppressive phenotype (CD11b^+^CD163^+^) monocytes among total CD45^+^ cells, indicating tumor-driven monocyte activation and polarization toward an M2-like immunosuppressive phenotype **(Fig. 2e and f)**. In contrast, direct coculture with CCHMC-DIPG-1 and CCHMC-DIPG-2 did not increase the proportion of CD11b^+^CD163^+^ monocytes, and their conditioned media had no observable effect on monocyte activation or polarization. These findings suggest that different DIPG cell lines exhibit variable capacities to recruit, activate, and polarize monocytes *in vitro*.

**Figure 2.**
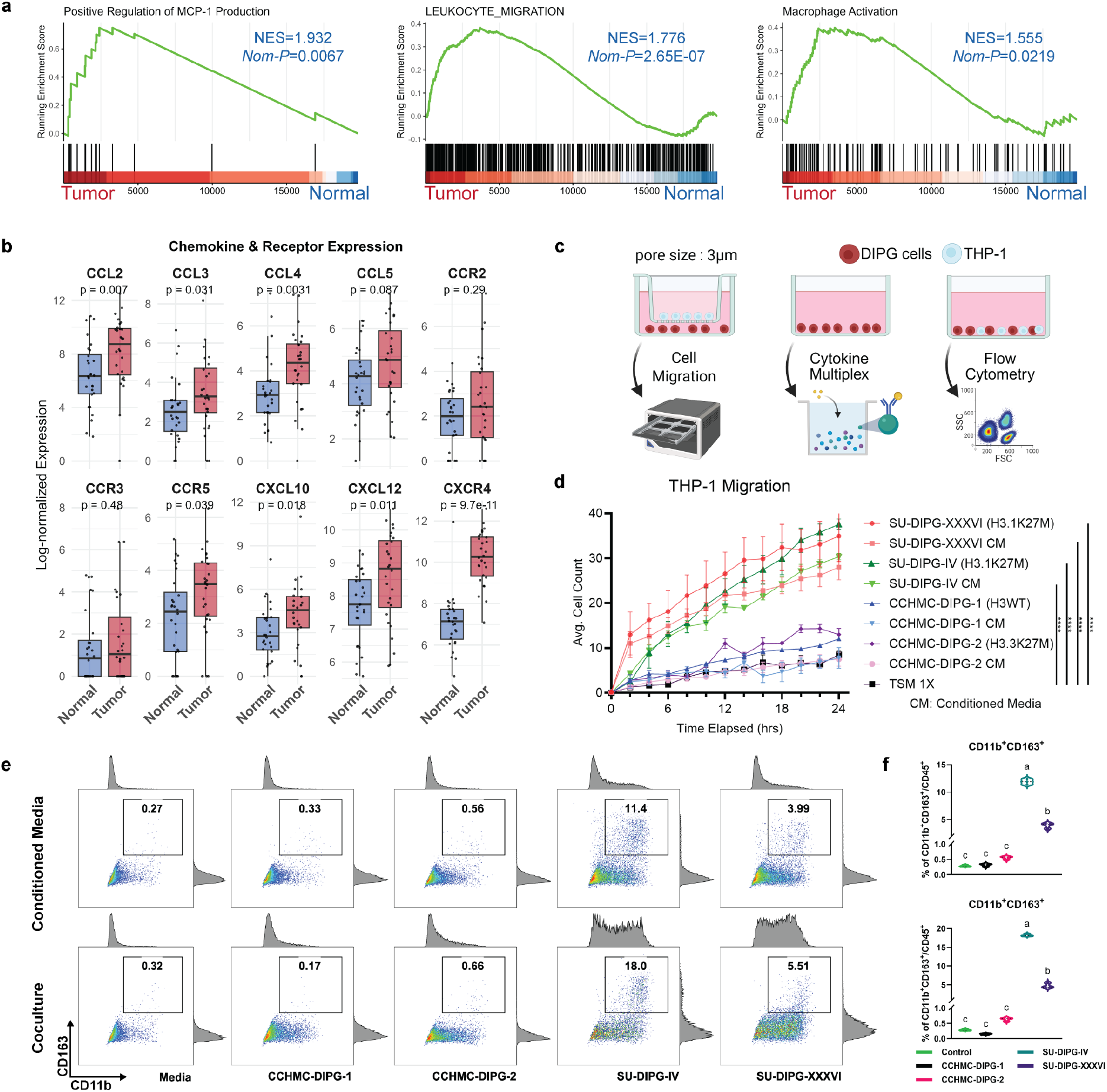
DIPG cells promote monocyte recruitment and activation *in vitro*. **(a)** Enriched positive regulation of MCP-1 production, leukocyte migration, and macrophage activation pathways in DIPG tumor samples through GSEA analysis. **(b)** Chemokine and chemokine receptor genes were universally upregulated in DIPG tumor tissue compared to matched normal brain tissue. **(c)** Schematic workflow of THP-1 and DIPG cells co-culture, transwell migration, multiplex cytokine assay, and flow cytometry (created with BioRender). **(d)** THP-1 cells exhibit increased migration toward SU-DIPG-IV and SU-DIPG-XXXVI cells, but not toward CCHMHC-DIPG-1 and CCHMC-DIPG-2. **(e)** Representative flow cytometry plots showing the proportion of CD11b^+^CD163^+^ cells out of total CD45^+^ cells, indicating polarization of M2-like immunosuppressive macrophages. **(f)** Flow cytometry results show variable capacities of DIPG cells to activate and polarize monocytes. Statistical difference was determined by two-way ANOVA; **** *p*< 0.0001; or One-way ANOVA with Tukey’s post-hoc test, groups with different letters are significantly different.

### Chemokine signatures in DIPG cells are lineage-dependent rather than driven by histone mutations

As noted previously, DIPG tumor cells exhibit broad molecular heterogeneity, including histone alterations and lineage identity, both of which may influence their capacity to modulate immune cells. To further investigate how DIPG cells drive monocyte migration and activation, and to define the molecular features underlying their variable capabilities, we analyzed cytokine and chemokine secretion using Luminex-based multiplex cytokine assay on 24-hour conditioned media from four DIPG cell lines **(Fig. 3a)**. Here, SU-DIPG-IV and SU-DIPG-XXXVI, which previously demonstrated a stronger capacity to recruit and polarize monocytes, exhibited markedly higher secretion levels of several key cytokine and chemokines, including CCL2, IL-10, and TGF-β1 **(Fig. 3b)**. In contrast, CCHMHC-DIPG-1 and CCHMC-DIPG-2 displayed a globally reduced secretion of these cytokines and chemokines. To validate these observations at a broader scale, we analyzed RNA-seq data from 46 pHGG cell lines available in the Childhood Cancer Model Atlas (CCMA), focusing on the expression of nine chemokines previously implicated in immune cell recruitment [67]. The analysis revealed marked heterogeneity in chemokine expression profiles across cell lines; however, this variability did not appear to follow any consistent pattern based on histone mutation status **(Fig. 3c)**. For further exploration, we applied single-sample gene set enrichment analysis (ssGSEA) to compute a chemokine expression score across pHGG cell lines. Although variability within each group was observed, no statistically significant differences were detected in chemokine scores **(Fig. 3d)** or in the expression of individual chemokines **(Supp. Fig. 2b)** across histone mutation subtypes.

**Figure 3.**
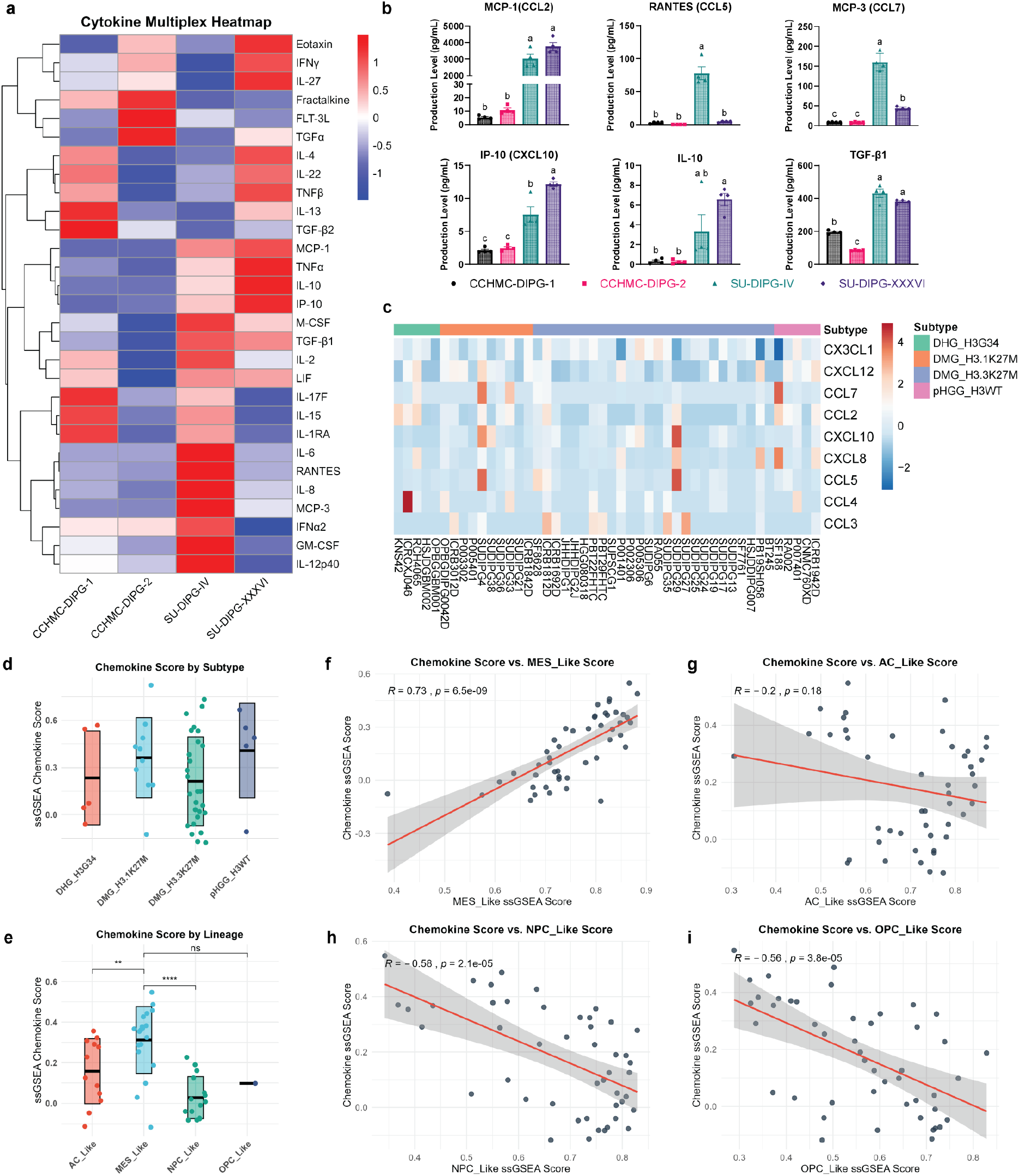
Chemokine expression differs across DIPG cell lines and correlates with intrinsic lineage states. **(a)** Multiplex cytokine assay shows distinct cytokine secretion profiles across DIPG cell lines. **(b)** Production level of key cytokines and chemokines in DIPG cell lines. **(c)** Heatmap of RNA-seq data from 46 pHGG cell lines reveals expression profiles of nine key chemokines. **(d)** ssGSEA chemokine expression scores across pHGG cell lines grouped by histone mutation subtypes. **(e)** MES-like cell lines showed higher chemokine scores compared to cell lines of other lineages. **(f)** Chemokine scores correlate with mesenchymal-like lineage program score in DIPG. **(g)** Chemokine scores are not related to astrocyte-like lineage program. **(h)** Chemokine scores negatively correlate with neuronal progenitor-like lineage program in DIPG. **(i)** Chemokine scores negatively correlate with oligodendrocyte precursor-like lineage programs in DIPG. Statistical difference was determined One-way ANOVA with Tukey’s post-hoc test, groups with different letters are significantly different, ** *p*<0.01, \*\*\*\**p*<0.001.

Recent studies showed that pediatric glioma and GBM tumor cells display distinct lineage states, including oligodendrocyte precursor-like (OPC-like), astrocyte-like (AC-like), neural progenitor-like (NPC-like), and mesenchymal-like (MES-like), which influence tumor cell behavior, therapeutic resistance, and TME interactions [1, 59, 79]. To evaluate the correlation between chemokine expression and intrinsic lineage state, we calculated lineage scores using ssGSEA in pHGG cell lines, based on selected marker gene sets for each lineage derived from Neftel et al., and evaluated their correlation with chemokine scores [59]. As the result suggested, chemokine scores were positively correlated with MES-like lineage scores (*R* = 0.73, *p* = 6.5e-09), indicating that tumor cells with mesenchymal features tend to express higher levels of chemokines **(Fig. 3e)**. In contrast, there was no significant correlation between chemokine expression and AC-like score (*R* = −0.2, *p* = 0.18) and significant negative correlation with OPC-like score (*R* = −0.56, *p* = 3.8e-05) and NPC-like score (*R* = −0.58, *p* = 2.1e-5) **(Fig. 3f-h)**. We then classified each pHGG cell line according to its dominant lineage score. Consistent with our correlation analysis, MES-like cell lines exhibited significantly higher chemokine scores than both AC-like and NPC-like cell lines **(Fig. 3i)**, as well as higher expression of several individual chemokines, such as *CCL2, CCL5*, and *CCL7* **(Supp. Fig. 2c)**. Although the OPC-like category contained only one cell line and thus could not be statistically compared, its chemokine score appeared lower than that of MES-like cell lines. Together, these data support that MES-like lineage state, rather than histone mutation status, is more strongly associated with enhanced chemokine production capacity among pHGG cell lines.

### Single-cell RNA sequencing analysis shows tumor-associated macrophage (TAM)–tumor cell crosstalk through multiple signaling pathways

To dissect the intratumoral heterogeneity and cell-cell communication at the single-cell resolution, we performed scRNA-seq on eight DIPG patient samples and integrated these data with publicly available pHGG scRNA-seq datasets (GSE162989, GSE184357, GSE131928), which encompass H3K27M-mutant, G34R/V-mutant, and H3 wildtype tumors [49, 48, 59]. Harmony was applied to correct for batch effects, enabling cross-study comparison [38]. After quality control, a total of 22,300 cells were retained for downstream analysis, encompassing H3WT, H3K27M, and G34R/V subtypes. Uniform manifold approximation and projection (UMAP) visualization and cell clustering revealed distinct cellular populations corresponding to malignant and non-malignant compartments **(Fig. 4a)**. We identified clusters representing subtypes of malignant cells, normal brain populations, and immune compartments, such as TAMs and tumor-infiltrating lymphocytes (TILs). Further stratification of the integrated scRNA-seq dataset by histone mutation subtype showed that all subtypes have a prominent proportion of TAMs within TME **(Fig. 4b)**. TAMs were consistently detected across all DIPG subtypes, although their relative abundances varied between patients, which may reflect both biological diversity and technical differences inherent to single-cell sample processing. Analysis of cellular composition across individuals showed substantial inter-patient heterogeneity, with variable proportions of both malignant and non-malignant populations **(Fig. 4c and d)**. The persistence of TAM composition across samples indicates that TAMs constitute a major component of the pHGG TME and play a central role in the immune landscape. Given the complexity of cellular composition in pHGG, we next applied CellChat analysis to construct a global map of intercellular communication and to better characterize the signaling networks shaping the TME [32]. Global signaling strength of incoming and outgoing interactions and the integrated communication network indicated that OPC-like, hypoxia-driven, and AC/MES-like tumor cells displayed high levels of both sending and receiving interactions **(Fig. 4e and f)**. In contrast, the non-malignant populations contributed relatively fewer overall signals, suggesting these populations might have a more specialized rather than a global communication role.

**Figure 4.**
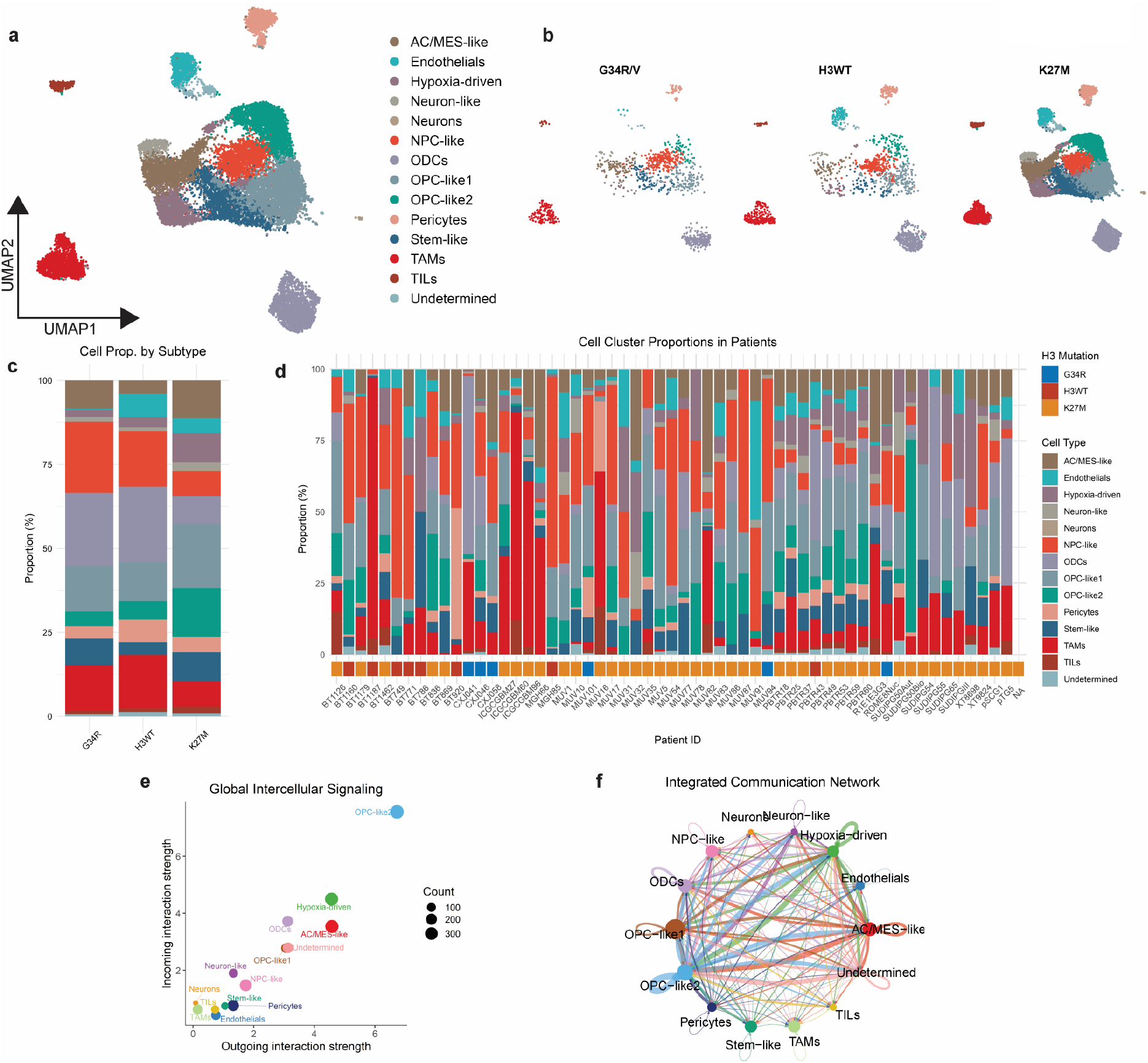
Single-cell transcriptomics landscape and intercellular communication networks in pediatric high-grade glioma. **(a)** UMAP visualization of all cells from pHGG samples, including K27M (n=43), G34R/V (n=6), and H3WT (n=8), color of cells indicates cell types. **(b)** UMAP of distribution of cells across different histone mutation subtypes of pHGG, color of cells indicates cell types. **(c)** Proportion of each annotated cell type within different subtypes of pHGG. **(d)** Proportion of each cell type in individual patients of pHGG. **(e)** Dotplot of the relative outgoing and incoming interaction strength of each cell type. **(f)** Integrated global cell communication network in pHGG.

Beyond overall communication intensity, we examined the specific signaling pathways with a focus on TAMs. In terms of outgoing signals, TAMs predominantly communicate with other populations through the secreted phosphoprotein 1 (SPP1) and progranulin (GRN) signaling pathways, consistent with previously reported roles of TAMs in brain tumors **(Supp. Fig. 3a)** [68, 80]. In addition, TAMs were also predicted to receive incoming signals from multiple cell types, most notably through amyloid precursor protein (APP), transforming growth factor-β (TGF-β), and semaphorin-3 (SEMA3) pathways, suggesting multiple potential axes of crosstalk that may regulate TAM function within the TME **(Supp. Fig. 3b)**. Among these pathways, TGF-β pathway has been well-recognized for driving an immunosuppressive macrophage phenotype, inducing PD-1 expression, and promoting the secretion of anti-inflammatory cytokines, such as IL-10 [20, 44, 83]. The expression of TGF-β in brain tumors has been extensively reported in multiple studies [45, 71]. In addition, TAMs were the sole recipient of SEMA3 signaling, which has been reported to function as a retention signal for macrophages by inhibiting their chemokine responsiveness [30, 75]. Collectively, SEMA3 and TGF-β pathways may contribute to retaining TAMs within the TME and promoting their polarization toward an M2-like immunosuppressive phenotype.

### The APP-CD74 signaling axis represents a prominent pathway targeting TAMs, despite the overall downregulation of APP in pHGG tumors

CellChat analysis predicted the APP-CD74 signaling axis as a prominent incoming cell-cell contact pathway in TAMs, although its role in DIPG has been minimally explored to date **(Fig. 5a-c)**. APP is a transmembrane glycoprotein broadly expressed in the central nervous system and its role in Alzheimer’s disease was originally explored [22]. CD74, also known as MHC class II in variant chain, functions as both membrane-bound receptor that transduce signals upon ligand binding and chaperone for MHC class II antigen presentation [63]. Visualization of gene expression on UMAP revealed that *APP* was broadly expressed across most cell populations within TME, whereas *CD74* was restricted to TAMs **(Fig. 5d and e)**.

**Figure 5.**
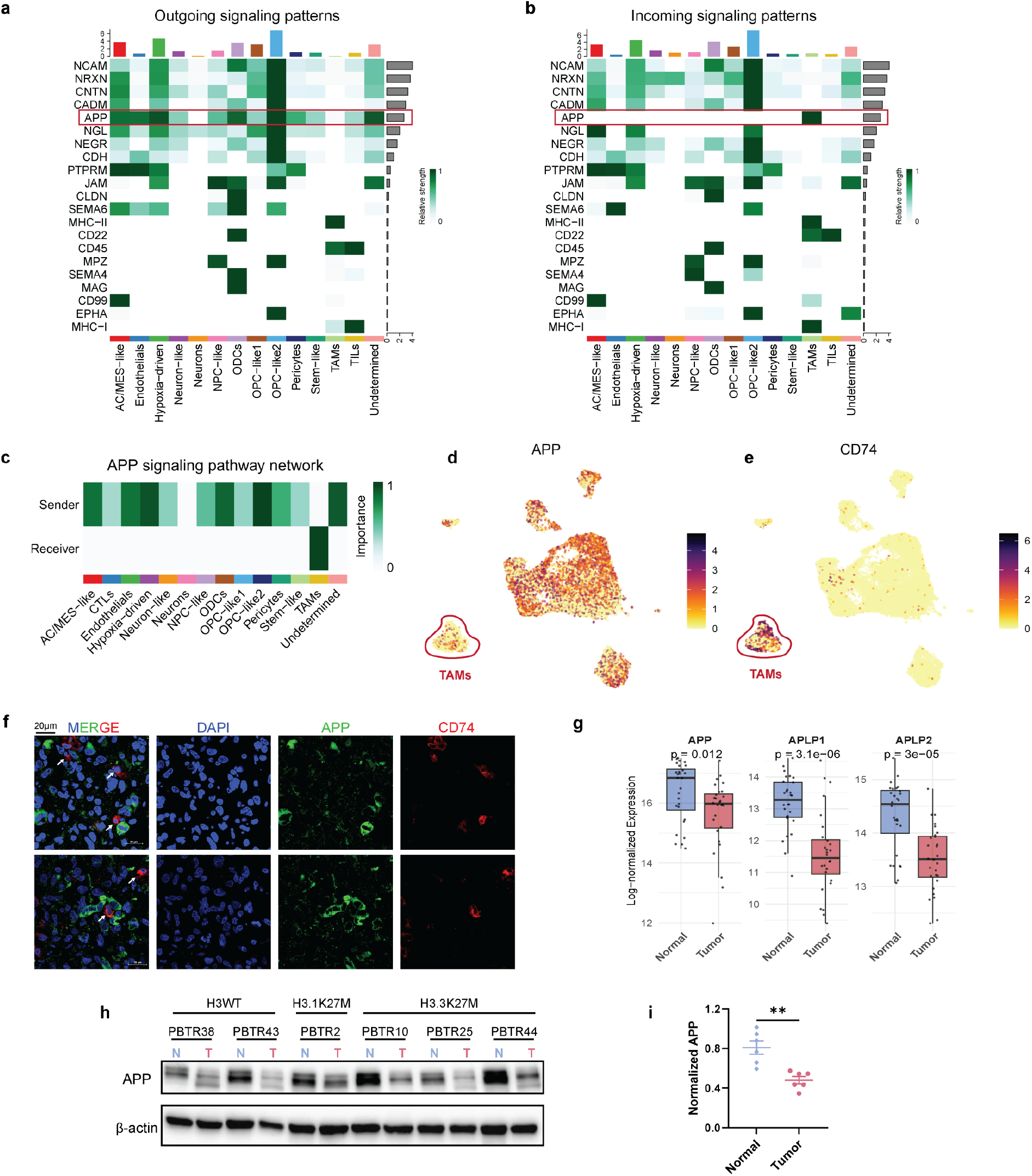
APP-CD74 emerges as a key signaling pathway in pHGG TME, while APP expression level is reduced in tumor tissues. **(a)** Outgoing signaling pathways with high relative strength. **(b)** Incoming signaling strength highlights TAMs as the only receiver of the APP-CD74 signaling pathway. **(c)** APP-CD74 signaling interactions in the TME of pHGG. **(d)** Visualization of APP expression on the UMAP projection plot. **(e)** Visualization of CD74 expression on the UMAP projection plot. **(f)** Immunofluorescence of DIPG tumor tissue sections demonstrates the spatial proximity of APP (Green) and CD74 (Red). **(g)** Reduced expression level of *APP*, and two other members of amyloid precursor protein family (*APLP1* and *APLP2*), in DIPG tumor tissues compared with matched normal brain tissues. **(h)** Western blot showing APP and β-actin (loading control) protein level in matched normal brain tissue (N) and tumor tissue (T) from six DIPG patients across three histone mutations. **(i)** Quantification of APP protein expression normalized to β-actin (APP/β-actin ratio). Data are presented as mean ± SEM. Statistical significance was determined by paired *t*-test, ** *p*< 0.01.

To validate the spatial colocalization of APP and CD74, we performed immunofluorescence co-staining of APP and CD74 in pHGG tumor sections. The results confirmed the spatial proximity between APP and CD74, supporting the ligand-receptor interaction within the TME **(Fig. 5f)**. Interestingly, comparative analysis suggests reduced *APP* expression level in tumor specimens from DIPG patients relative to paired normal brain tissues. Consistently, bulk RNA-seq analysis demonstrated that *APP* expression was significantly lower in tumor regions than in normal brain tissue, along with decreased expression of *APLP1* and *APLP2*, the other members of the APP protein family **(Fig. 5g)**. The reduced production of APP in DIPG tumor tissue was further confirmed by Western blot analysis **(Fig. 5h and i)**. Collectively, these data support the presence of APP-CD74 signaling directed toward TAMs in pHGG, suggesting a functional role for this signaling axis in modulating macrophage behavior.

### APP stimulation induces a pro-inflammatory response in macrophages

To functionally validate the effect of APP on macrophage polarization and function, we treated THP-1 derived macrophages with recombinant human APP and assessed the downstream transcriptional changes and cytokine secretion. The APP treatment induced substantial transcriptional reprogramming in THP-1 derived macrophages, as revealed by RNA-seq **(Fig. 6a)**. Differential expression analysis identified 340 upregulated and 110 downregulated genes (adjusted *p* value <0.05, |log2FC|>1). Of particular interest, APP stimulation markedly induced the expression of multiple pro-inflammatory markers, including interferon-stimulated genes (*IFIT1* and *IFIT2*) and inflammatory cytokines (*IL1B, CCL2*, and *CXCL10*). In parallel, APP stimulation inhibited the expression of canonical M2-associated markers, including *CD163, MRC1* (CD206), and *ARG2* **(Fig. 6a)**. Consistently, Gene Set Enrichment Analysis (GSEA) of Hallmark pathways revealed strong enrichment of inflammatory pathways, including inflammatory response (NES = 1.500, *adj*.*p* =5.24e-3), interferon alpha response (NES = 1.806, *adj*.*p* = 5.28e-05), interferon gamma response (NES = 1.730, *adj*.*p* = 7.70e-05), and TNF alpha signaling via NF-κB (NES = 1.622, *adj*.*p* = 6.01e-04) **(Fig. 6b)**. Together, this reciprocal pattern of induction of inflammatory response suppression and M2-like markers suggests a transcriptional shift toward a pro-inflammatory phenotype.

**Figure 6.**
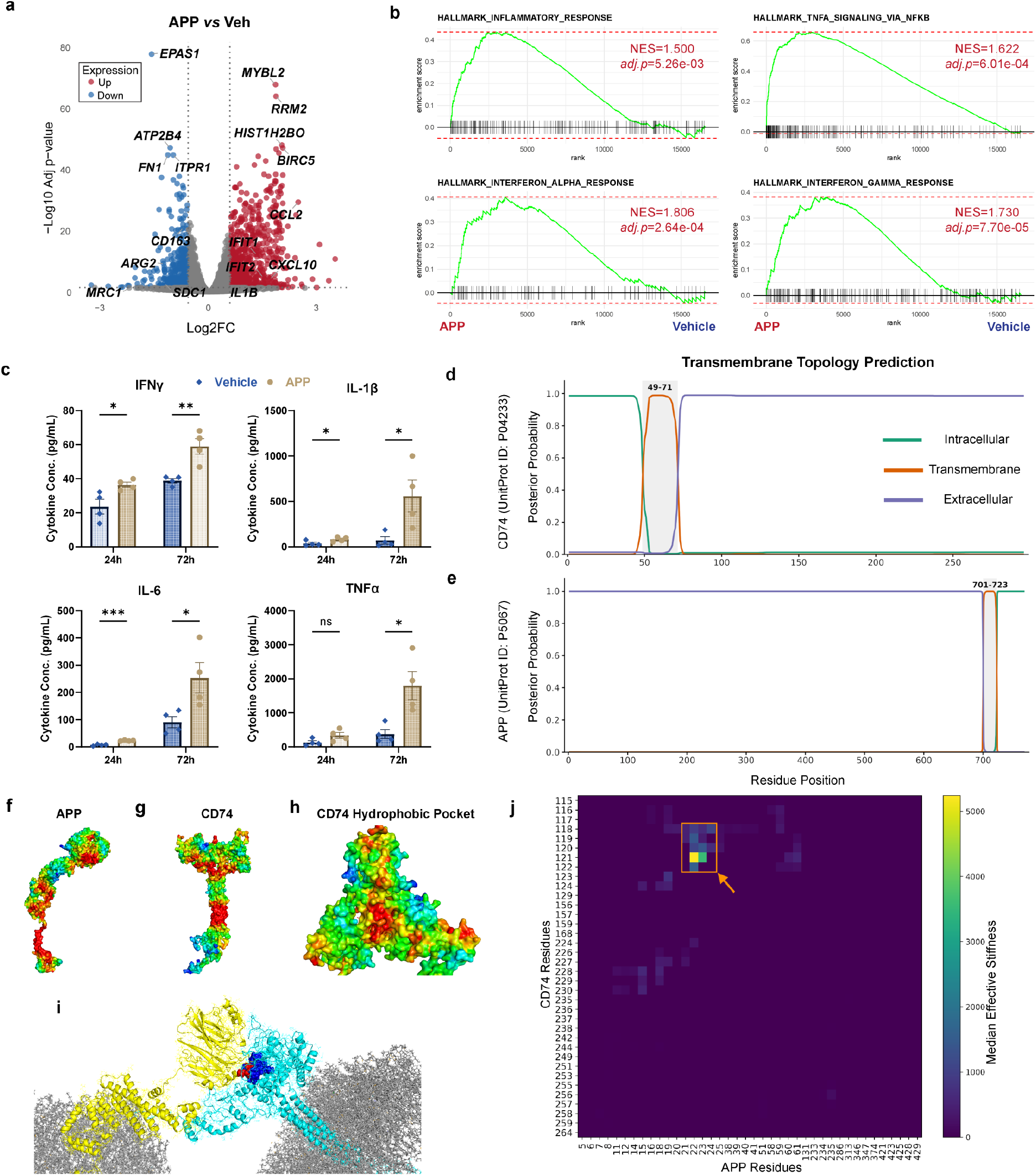
APP stimulation induces proinflammatory response in THP-1-derived macrophages and structural modeling identifies APP-CD74 interaction interface. **(a)** Volcano plot of RNA-seq data showing differentially expressed genes in THP-1 derived macrophages treated with recombinant APP protein compared to vehicle control. **(b)** GSEA of RNA-seq data from APP-treated THP-1-derived macrophages reveal significant enrichment of Hallmark proinflammatory-related pathways. **(c)** Multiplex cytokine assay of conditioned media from APP-treated THP-1-derived macrophages (24 and 72 hours) shows increased secretion levels of IFNγ, IL-1β, IL-6, and TNFα compared with vehicle control. Data are presented as mean ± SEM. Statistical significance was calculated with unpaired student *t*-test, *p<0.05; **p<0.01; ***p<0.001; ns, not significant. **(d)** TMHMM-predicted transmembrane regions of APP. **(e)** TMHMM-predicted transmembrane regions of CD74. **(f)** Surface hydrophobic patch mapping of APP. The color gradient from blue to red represents increasing relative hydrophobicity, with blue indicating low hydrophobicity and red indicating high hydrophobicity. **(g)** Surface hydrophobic patch mapping of CD74. The color gradient from blue to red represents increasing relative hydrophobicity, with blue indicating low hydrophobicity and red indicating high hydrophobicity. **(h)** Central hydrophobic pocket of the CD74 oligomer. The color gradient from blue to red represents increasing relative hydrophobicity, with blue indicating low hydrophobicity and red indicating high hydrophobicity. **(i)** Membrane-restrained Gaussian network model analysis showing median effective stiffness of residue pairs at the APP-CD74 interface. The color gradient from blue to yellow represents increasing median effective stiffness, with blue indicating low median effective stiffness and red indicating high median effective stiffness. **(j)** Docked APP-CD74 complex showing interaction of the APP N-terminal region with the central hydrophobic pocket of CD74 in the membrane-associated arrangement. Membrane is shown in grey, APP is shown in yellow, CD74 oligomer is shown in cyan, APP residue 19-24 is shown in red spheres, and CD74 residue 118-122 is shown in blue spheres.

To determine whether these transcriptional changes translate into altered cytokine secretion, we measured cytokine levels in culture supernatants at 24 and 72 hours following APP treatment. APP stimulation induced a time-dependent increase in multiple pro-inflammatory cytokines. Consistent with the RNA-seq results, APP treatment significantly elevated the secretion of pro-inflammatory cytokines, including IFN-γ, TNF-α, IL-1β, and IL-6 **(Fig. 6c)**. Meanwhile, APP treatment also significantly increased the secretion of multiple myeloid growth factors and chemoattractants that have been shown to enhance anti-tumor immunity in preclinical glioma models **(Supp. Fig. 4)** [16, 61, 62].

Together, these results indicate that APP stimulation drives macrophages toward a pro-inflammatory, M1-like activation state characterized by activation of interferon signaling pathways, upregulation of pro-inflammatory gene expression, and enhanced secretion of inflammatory cytokines. This APP-induced pro-inflammatory phenotype may contribute to the establishment of an immunologically active niche within the TME and suggests that restoring APP signaling in tumor regions could represent a potential avenue for immunotherapeutic intervention.

### Structural modeling reveals a high-stiffness binding interface between APP and CD74

To investigate the structural basis of the APP–CD74 interaction, we performed topology prediction, hydrophobic surface mapping, protein–protein docking analyses, and Gaussian network modeling of membrane-bound APP and CD74. TMHMM-based topology prediction identified a single high-confidence transmembrane segment in APP spanning residues 701-723 and in CD74 spanning residues 49-71 **(Fig. 6a and b)**. Hydrophobic surface patch mapping further revealed that both proteins contain multiple nonpolar regions that may contribute to membrane association and intermolecular recognition. In APP, hydrophobic segments were detected at residues 5-17, 343-349, 357-379, 481-498, 603-614, 643-

649, 654-676, 686-745, and 757-761 **(Fig. 6c)**. Notably, the broad C-terminal hydrophobic region (686-745) fully encompasses the predicted transmembrane helix (701-723) of APP, consistent with a membrane-embedded C-terminal domain architecture. In CD74, hydrophobic patches were identified at residues 49-72, 95-124, 156-165, 181-189, 204-225, and 284-297 **(Fig. 6d)**. The hydrophobic segment 49-72 overlaps with the predicted CD74 transmembrane region (49-71), consistent with the experimentally established membrane anchoring role of N-terminal [15, 63]. In addition, CD74 also displayed a large hydrophobic pocket within residues 110-124, located within the central region of the CD74 oligomer **(Fig. 6e)**. Because of the spatial accessibility and nonpolar residue enrichment, it represented a plausible interaction site for the APP N-terminal region and was selected as the docking interface for protein-protein docking analysis with the APP N-terminal region.

Membrane-restrained Gaussian Network Modeling (GNM) analysis of the docked APP-CD74 complex was performed to identify the interfacial regions of high mechanical coupling. The strongest coupling was localized to a narrow CD74 segment centered on residues 118-122, which interacts with APP residues 19-24 **(Fig. 6f and g)**. Another coupling was observed at residues 116-124 and 224-230 of CD74, suggesting a stabilizing role of these regions at the binding interface **(Fig. 6f)**. The strong enrichment of high-stiffness couplings within the central hydrophobic pocket of CD74 oligomer indicates that this segment acts as a conformational constraint between APP and CD74. Together, these findings identify CD74 residues 118-122 and APP residues 19-24 as mechanically critical determinants of the APP-CD74 interface, providing a structurally defined target for potential therapeutic interventions.

## DISCUSSION

In this study, we present a comprehensive characterization of the TME in DIPG, one of the most aggressive types of pediatric brain tumors, through integrated multi-omics analysis. Our results indicate that DIPG maintains a myeloid-dominant environment in which TAMs are actively recruited and functionally shaped by paracrine signals from tumor cells, particularly mesenchymal-like tumor cells. CellChat analysis of scRNA-seq data identified the APP-CD74 signaling axis as a key intercellular communication pathway within the pHGG TME. Notably, DIPG tumor tissues show decreased *APP* expression, which may contribute to the immunosuppressive phenotype of TAMs, given that APP drives pro-inflammatory programs in macrophages. Collectively, these findings improve our understanding of the sophisticated interplay between tumor and immune compartments in the pHGG microenvironment and identify the APP–CD74 signaling axis as a potential therapeutic target for remodeling the TME. Our bulk RNA-seq analysis confirmed the previously reported enrichment of myeloid cells in DIPG and further revealed broad upregulation of genes involved in MHC antigen presentation pathways [14, 46, 56]. The discrepancy between the upregulation of antigen presentation machinery, characterized by MHC expression, and the paucity of cytotoxic T lymphocytes likely reflects a dysfunctional immune interface rather than a functional anti-tumor immune response. This interpretation is further supported by recent findings suggesting that MHCII expression on blood-borne myeloid cells is required for effective anti-tumor T cell responses, whereas antigen presentation by TAMs may instead contribute to the terminal exhaustion of T cells derived from a progenitor exhausted state [37, 73].

Our Transwell and flow cytometry assays indicated that DIPG cells can recruit and polarize monocytes, although this capacity varies across cell lines. Further bioinformatics analysis revealed that the chemokine production capacity of DIPG cells is significantly associated with the MES-like lineage identity (R = 0.73, *p* = 6.5e-09) rather than histone mutation status. These findings established a crucial role of MES-like glioma cells in promoting a myeloid-dominant immune environment in pHGG and in polarizing TAMs toward a non-inflammatory state. The elevated chemokine expression profile observed in MES-like glioma cells may be driven by activation of NF-κB and STAT3 signaling pathways, key transcriptional regulators of *CCL2* and *CCL5* expression [7, 8, 31, 74]. Intriguingly, a recent study in GBM has demonstrated that M2-like TAMs promote mesenchymal transition in tumor cells, suggesting a self-reinforcing feedback loop between MES-like tumor cells and TAMs within the brain tumor microenvironment: MES-like tumor cells recruit and polarize peripheral macrophages toward an M2-like state, which in turn stabilizes the mesenchymal phenotype of tumor cells [35, 79, 82].

Furthermore, CellChat analysis highlights the APP-CD74 axis as a noteworthy cell-cell contact pathway to TAMs. APP was first described as a direct binding ligand of CD74 in the context of Alzheimer’s disease that suppresses amyloid-β (Aβ) production, while we showed the spatial proximity between APP and CD74-expressing cells in the DIPG tumor sites in this study [52]. Direct binding between APP and CD74 proteins has been shown in multiple studies using immunoprecipitation assays [47, 51]. To elucidate the structural interface mediating APP-CD74 interaction in a physiologically relevant membrane-bound context, we performed protein-protein docking with GNM analysis. By integrating the lipid bilayers of cell membrane, this framework identified the contact regions between APP and CD74, as well as the residues with elevated binding energies. Topology prediction and hydrophobic surface mapping depicted the membrane architecture of both proteins and informed the selection of binding interfaces. GNM analysis of the docked complex revealed the high-stiffness binding sites at the APP-CD74 interface within APP residues 19-24 and CD74 residues 118-122, which are embedded in the hydrophobic pocket of the CD74 oligomer. These computational findings provide structural evidence supporting the physical interaction between APP and CD74 that was observed in previous experiments, and highlighting specific binding residues as actionable targets for treatment [52].

Transcriptomic analysis and cytokine assays revealed that recombinant APP stimulation promotes pro-inflammatory polarization and inflammatory cytokine secretion in THP-1–derived macrophages, consistent with the canonical role of the CD74 receptor [63]. Combined with the reduced *APP* expression observed in DIPG tumor tissues at both RNA and protein levels, *APP* downregulation may attenuate the pro-inflammatory activation of TAMs, thus contributing to the immunosuppressive TME in pHGG. Integrating these findings with prior work extends the pathological relevance of APP from neurodegenerative disorders to brain tumors [18, 21, 43]. The abnormal processing of APP leads to neurotoxic amyloid-β peptide accumulation, which has been shown to modulate microglia function and prime the neuroinflammatory milieu of the central nervous system (CNS) [19, 22]. Accordingly, APP-mediated immune regulation may represent a pathological progression in various CNS diseases. A recent study in GBM reported that APP inhibits macrophage phagocytosis via CD74, implicating an immunosuppressive role of APP [51]. This discrepancy likely reflects the difference between the phagocytosis activity of macrophages in response to APP knockout in GBM cells and the transcriptional consequences of APP stimulation, which was not examined in the GBM study.

In summary, this study provides insights into the role of TAMs in pHGG TME and evidence to support that MES-like lineage tumor cells govern the recruitment of myeloid cells. It also identifies the APP-CD74 pathway as a functionally active immunoregulatory pathway in pHGG. The reduction of APP in tumor sites and the pro-inflammatory effect of APP treatment on macrophages suggest that diminished *APP* expression contributes to the immunosuppressive phenotypes of TAMs that are related to the poor prognosis of pHGG patients. These discoveries also expand our current understanding of TAMs in pediatric brain tumors and provide a rationale for potential treatment targeting the APP-CD74 axis. The high-stiffness interfacial region in the APP-CD74 interaction offers the possibility of developing peptides capable of engaging the CD74 receptor and reprogramming TAMs towards pro-inflammatory phenotypes with anti-tumor capacity.

### Limitation of Study

The bulk RNA-seq and scRNA-seq were performed on post-mortem autopsy tissues from patients, which might not fully recapitulate the TME of DIPG at earlier stages of tumor progression. In addition, patients in this cohort received various treatments that could influence the immune microenvironment or alter the phenotypes of TAMs, thereby introducing potential confounding effects in the interpretation of transcriptomic profiles, including those related to immune compartments. Future studies incorporating biopsy specimens or treatment-stratified cohorts will be important to disentangle treatment-related effects arising from intrinsic tumor and immune biology.

In addition, the GNM-based analysis identifies potential interfacial regions of the APP–CD74 complex; however, the model does not fully capture the dynamic, membrane-constrained conditions in which this interaction occurs. Further experimental validation, including mutagenesis of key residues in APP and CD74 and co-immunoprecipitation assays, will be necessary to confirm these findings in a cellular context. Moreover, although THP-1 cells and THP-1–derived macrophages are widely used as models of human monocytes/macrophages, they do not fully recapitulate the phenotypic and functional complexity of TAMs and may lack physiological relevance. Future studies employing primary human systems, such as peripheral blood–derived monocytes and patient-derived organoids, in conjunction with syngeneic preclinical mouse models, will be important to validate the translational significance of these findings.

## ETHICS STATEMENTS

## Acknowledgements

We thank the children and families who have generously donated tumor tissue for their invaluable contribution to this research. We acknowledge The International DIPG/DMG Registry Operations team for providing tissue samples. The authors also thank Gene Expression Core of Cincinnati Children’s Hospital Medical Center (CCHMC), Steve and Cindy Rasmussen Institute for Genomic Medicine (SCRIGM), Animal Resources Core (ARC), Flow Cytometry Core, Histopathology Core, and Microscopy Core at Nationwide Children’s Hospital (NCH).

## Author contributions

Conceptualization: Z.W., A.K., R.D.; Methodology: Z.W., A.K.; Formal analysis and investigation: Z.W., A.K., B.U., A.M.I., H.H.P., K.K.; Patient Identification and consent: M.F.; Writing-original draft preparation: Z.W.; Writing – review and editing: All authors; Funding acquisition: Z.W., R.D.; Supervision: R.D.

## Data Availability

Bulk RNA sequencing and single-cell RNA sequencing data will be deposited in European Genome-phenome Archive (EGA) and will be available upon request.

## Figure Legends

**Supplementary Figure 1.**
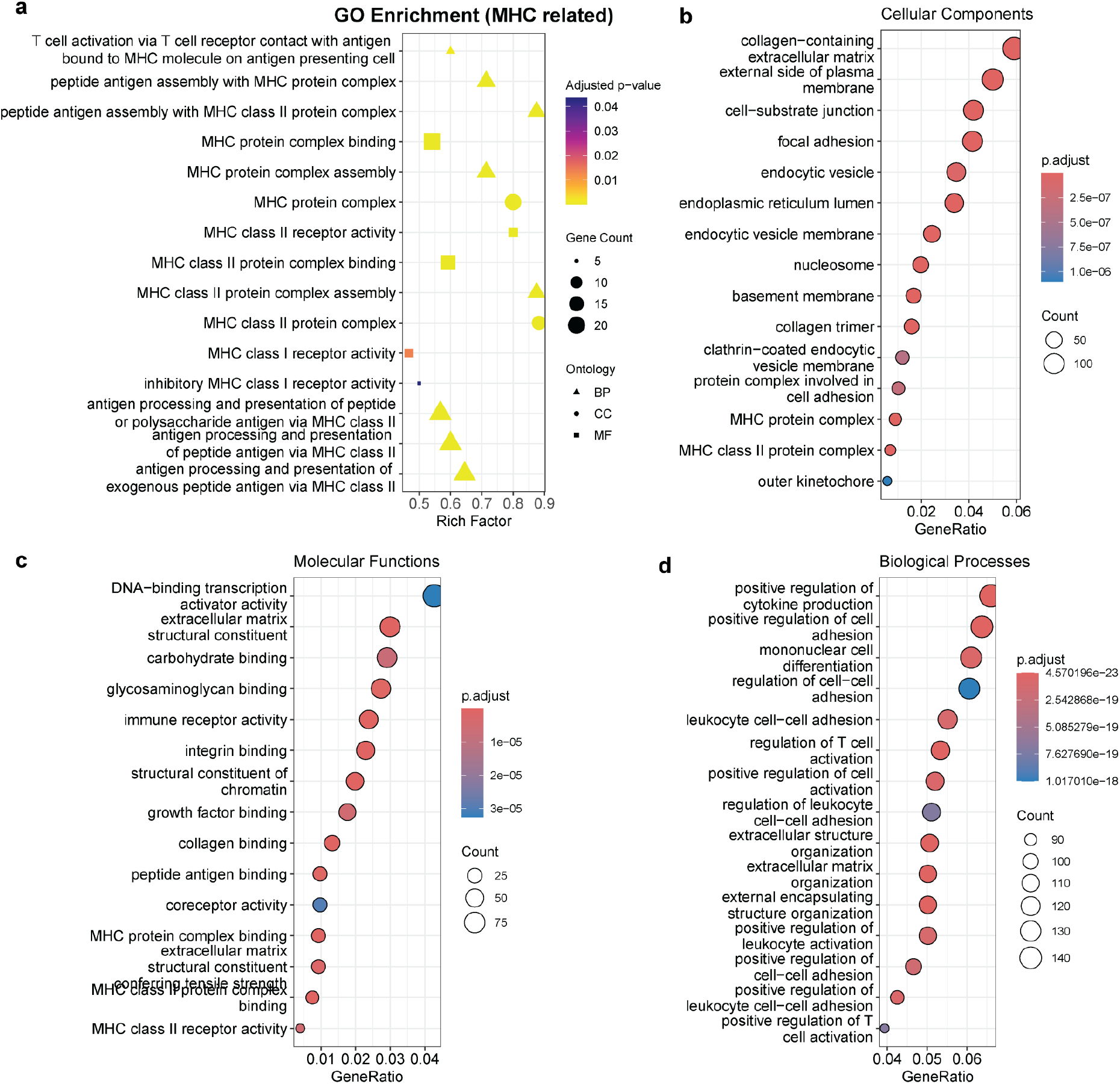
Summary of GO pathway enrichment analysis results. **(a)** MHC-related pathways among enriched Gene Ontology terms of upregulated genes in tumor specimens of DIPG patients. **(b)** Top 15 enrichment pathways within the cellular component (CC) category. **(c)** Top 15 enrichment pathways within the molecular feature (MF) category. **(d)** Top 15 enrichment pathways within the biological process (BP) category.

**Supplementary Figure 2.**
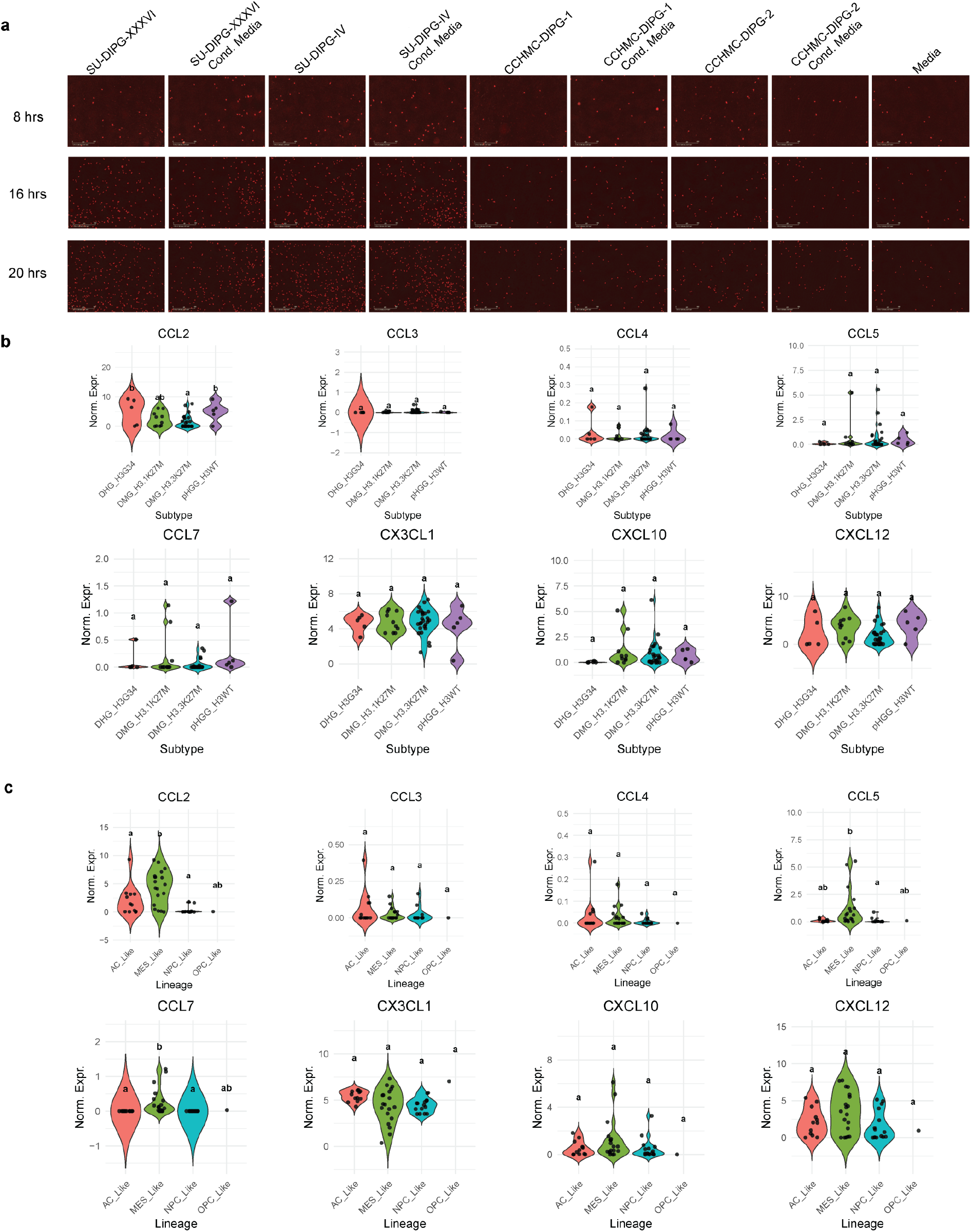
THP-1 monocyte migration and cytokine expression of 46 pHGG cell lines. **(a)** Representative Incucyte live cell images of transwell migration assay. THP-1 cells were stained with CellTracker CMRA (red). **(b)** Expression level of key chemokines in pHGG cell lines by histone mutation subtypes. **(c)** Expression level of key chemokines in pHGG cell lines by predominant cell lineage states.

**Supplementary Figure 3.**
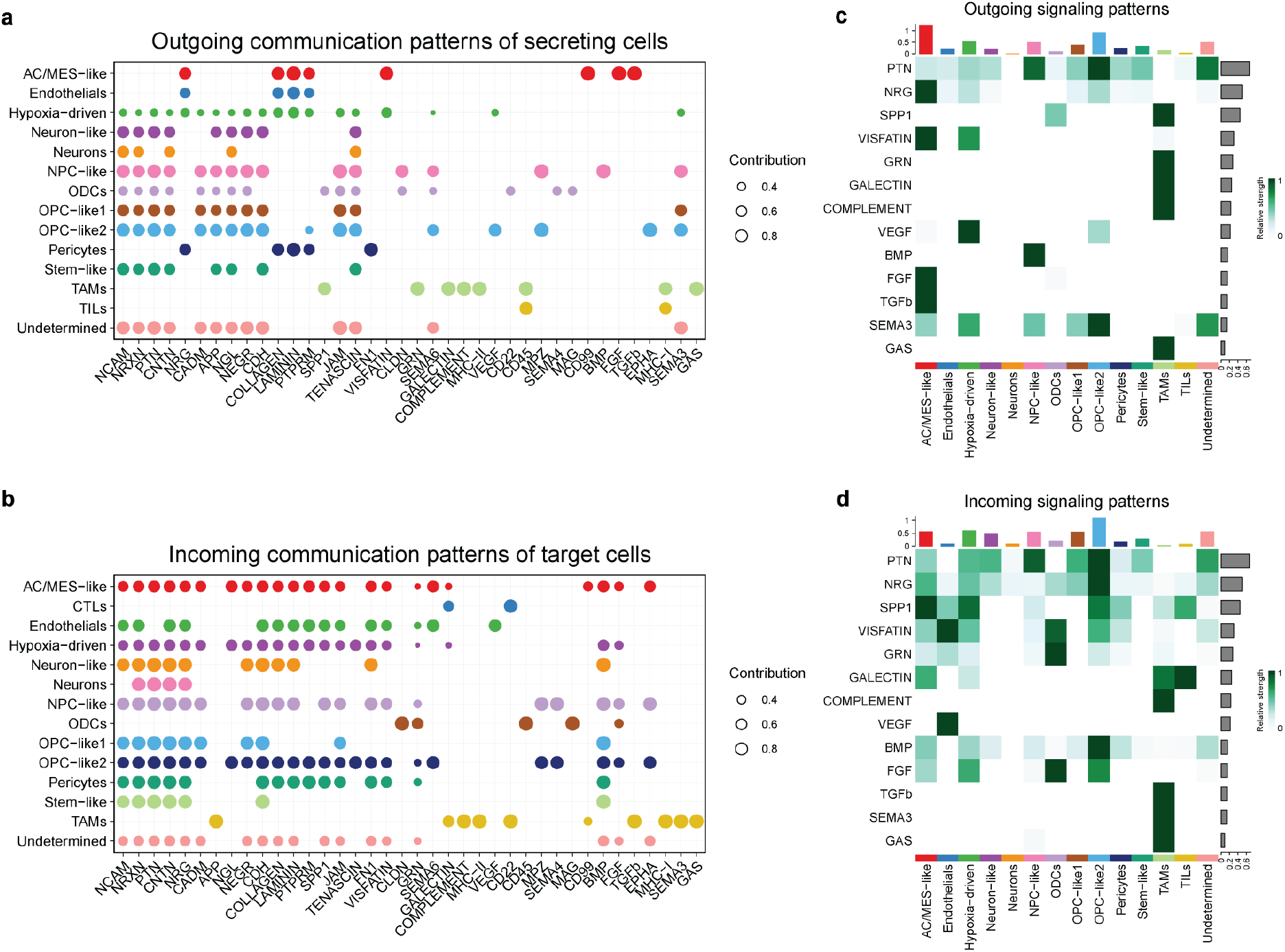
CellChat analysis of intercellular communication between distinct cell compartments in the pHGG microenvironment. **(a)** Outgoing signaling patterns of the global cell communication network. **(b)** Incoming signaling patterns of the global communication network. **(c)** Outgoing signaling patterns of the secreted signaling network. **(d)** Incoming signaling patterns of the secreted signaling network.

**Supplementary Figure 4.**
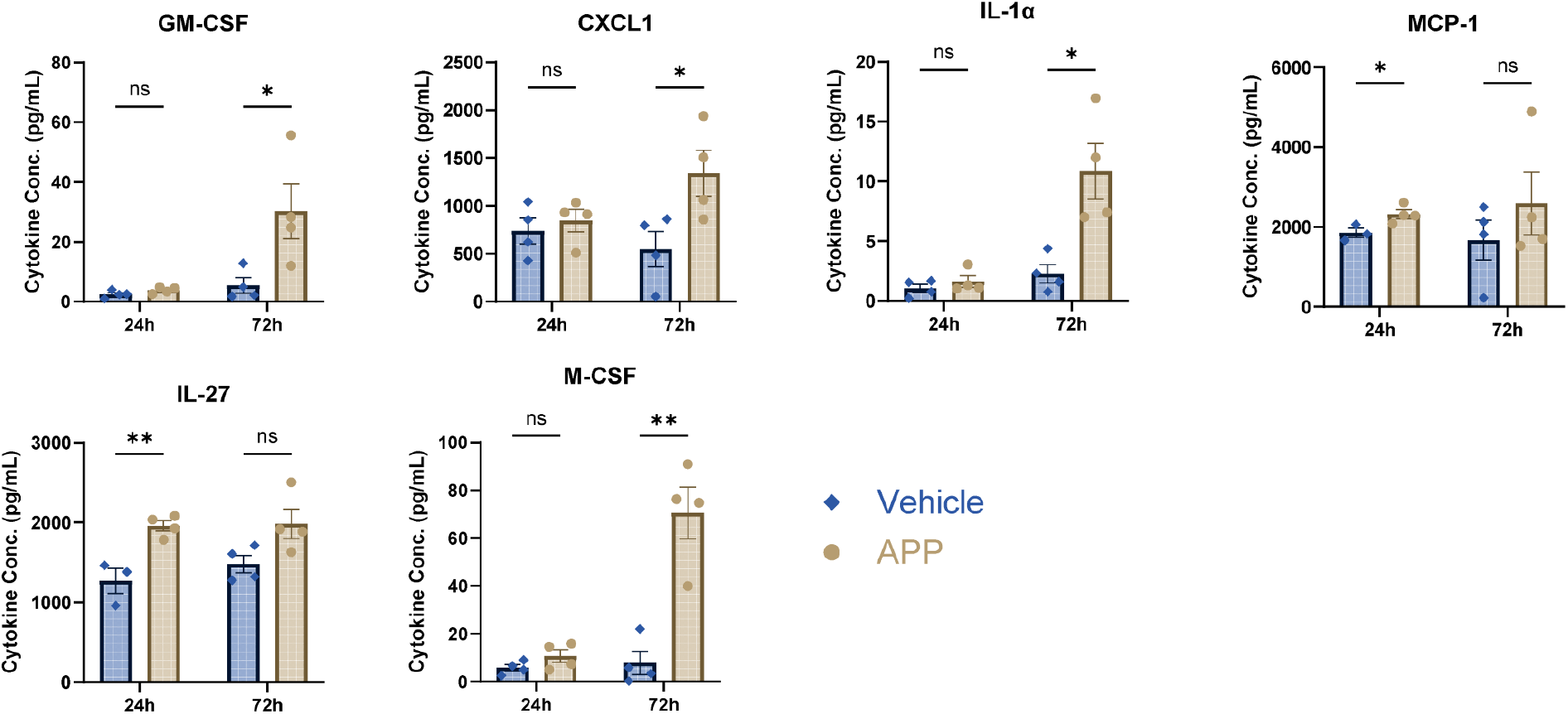
Cytokines and chemokines secretion by THP-1-derived macrophage upon APP stimulation. Multiplex cytokine assay of conditioned media from APP-treated THP-1-derived macrophages (24 and 72 hours) indicates the secretion level of GM-CSF, CXCL1, IL-1α, MCP-1, IL-27, and M-CSF compared to vehicle control. Data are presented as mean ± SEM. Statistical significance was calculated with unpaired student t-test, *p<0.05; **p<0.01; ns, not significant.

